# RFIDeep: Unfolding the Potential of Deep Learning for Radio-Frequency Identification

**DOI:** 10.1101/2023.03.25.534127

**Authors:** Gaël Bardon, Robin Cristofari, Alexander Winterl, Téo Barracho, Marine Benoiste, Claire Ceresa, Nicolas Chatelain, Julien Courtecuisse, Flávia A.N. Fernandes, Michel Gauthier-Clerc, Jean-Paul Gendner, Yves Handrich, Aymeric Houstin, Adélie Krellenstein, Nicolas Lecomte, Charles-Edouard Salmon, Emiliano Trucchi, Benoit Vallas, Emily M. Wong, Daniel P. Zitterbart, Céline Le Bohec

**Affiliations:** Centre Scientifique de Monaco, Département de Biologie Polaire, Monaco, Principality of Monaco; Université de Strasbourg, CNRS, IPHC UMR 7178, Strasbourg, France; University of Turku, Department of Biology, Turku, Finland; Friedrich-Alexander-University Erlangen-Nürnberg, Department of Physics, Erlangen, Germany; University of Moncton, Canada Research Chair in Polar and Boreal Ecology and Centre d’Études Nordiques, Department of Biology, Moncton, NB, Canada; Marche Polytechnic University, Department of Life and Environmental Sciences, Ancona, Italy; Université de Genève, Faculté des Sciences, Genève, Switzerland; Woods Hole Oceanographic Institution, Applied Ocean Physics and Engineering Department, Woods Hole, MA, USA; Beefutures, Nantes, France; Terres australes et antarctiques françaises, Saint-Pierre, La Réunion, France; Stanford University, Stanford, CA, USA

**Keywords:** Artificial intelligence, behaviour, machine learning, RFID, wildlife monitoring

## Abstract

1. Automatic monitoring of wildlife is becoming a critical tool in the field of ecology. In particular, Radio-Frequency IDentification (RFID) is now a widespread technology to assess the phenology, breeding, and survival of many species. While RFID produces massive datasets, no established fast and accurate methods are yet available for this type of data processing. Deep learning approaches have been used to overcome similar problems in other scientific fields and hence might hold the potential to overcome these analytical challenges and unlock the full potential of RFID studies.
2. We present a deep learning workflow, coined “RFIDeep”, to derive ecological features, such as breeding status and outcome, from RFID mark-recapture data. To demonstrate the performance of RFIDeep with complex datasets, we used a long-term automatic monitoring of a long-lived seabird that breeds in densely packed colonies, hence with many daily entries and exits.
3. To determine individual breeding status and phenology and for each breeding season, we first developed a one-dimensional convolution neural network (1D-CNN) architecture. Second, to account for variance in breeding phenology and technical limitations of field data acquisition, we built a new data augmentation step mimicking a shift in breeding dates and missing RFID detections, a common issue with RFIDs. Third, to identify the segments of the breeding activity used during classification, we also included a visualisation tool, which allows users to understand what is usually considered a “black box” step of deep learning. With these three steps, we achieved a high accuracy for all breeding parameters: breeding status accuracy = 96.3%; phenological accuracy = 86.9%; breeding success accuracy = 97.3%.
4. RFIDeep has unfolded the potential of artificial intelligence for tracking changes in animal populations, multiplying the benefit of automated mark-recapture monitoring of undisturbed wildlife populations. RFIDeep is an open source code to facilitate the use, adaptation, or enhancement of RFID data in a wide variety of species. In addition to a tremendous time saving for analyzing these large datasets, our study shows the capacities of CNN models to autonomously detect ecologically meaningful patterns in data through visualisation techniques, which are seldom used in ecology.

## 1 Introduction

Electronic monitoring systems have been widely used over the past two decades to better understand animal populations without human disturbance (Fagerstone & Johns, 1987; Schooley *et al*., 1993). Radio-frequency identification (RFID) technology allows the monitoring of uniquely identified individuals and automated recording of the presence of tagged individuals at chosen locations (Gibbons & Andrews, 2004). By placing RFID antennas along animal paths at perches or narrow entries of the breeding site (Gendner *et al*., 2005; Bonter & Bridge, 2011), individual survival and breeding rates as well as behaviour and locations can be precisely estimated in e.g. the classical capture-mark-recapture framework (Le Bohec *et al*., 2008). While RFID technology allows the recording of vast amounts of data, it also creates new challenges for data treatment, even if the data structure itself is rather simple (i.e., id, date and time, and location for each detection) (Iserbyt *et al*., 2018). Because RFID data are not directly linked with biological parameters, one of the classic approaches is human expert interpretation (Descamps *et al*., 2002; Afanasyev *et al*., 2015). Even today, most of the information extraction and ecological interpretation from such detection data is done manually, although this remains extremely time-consuming and potentially biased by human interpretation. Additionally, the difficulty in manually processing potentially large numbers of detection data is increased by the possibility of missing detections (Hughes *et al*., 2021).

A solution to these challenges may lie in automated data processing that could mimic the behaviour of an expert analyst. Artificial intelligence has been the focus of intense methodological effort in ecology: it has been used to process various sources of data including imagery or passive and active acoustic data, and to detect, classify, localise, identify, estimate, and predict at every biological scale, from individuals to ecosystems (Christin *et al*., 2019; Pichler & Hartig, 2022). Among artificial intelligence methods, deep learning has a wide and promising scope but often lacks approachable workflows for ecologists. Convolutional Neural Networks (CNN) have been initially developed for image content classification (Krizhevsky *et al*., 2017), but have also been used for classifying signals (Hinton *et al*., 2012) such as human activity classification (Mutegeki & Han, 2020), birds vocalisation classification (Kahl *et al*., 2021) or marine mammal detections (Shiu *et al*., 2020). Yet, CNN capacities remain unexplored in numerous fields such as RFID data processing.

Recent efforts have been made to automatically infer behavioural patterns from various types of biologgers through AI (Fannjiang *et al*., 2019; Wang, 2019): for instance, accelerometers have shown promising capacities to detect food-catching events (Brisson-Curadeau *et al*., 2021) or activity classification (Sakamoto *et al*., 2009; Jeantet *et al*., 2021). RFIDeep builds upon these efforts to address the specific nature of RFID data. While active biologgers record rich, multidimensional data, their record time is limited because of the required trade-off between miniaturisation, storage capacity, power consumption and impact on wildlife (Bodey *et al*., 2018). In contrast, passive, battery-less RFID tags have no demonstrable impact on an animal’s behaviour and function throughout the individual’s lifetime: but the trade-off is that although the tag is attached to the animal, detection only occurs at one or more fixed points (the active antennas), thereby offering a very narrow observation window. This creates a rather unique data structure with specific challenges for interpretation. RFID technology is also exposed to two major constraints because of the impossibility to detect multiple tags at the same time with a single antenna and the impossibility to install several antennas at the same place, due to electromagnetic interference. By increasing probability of missing detections, the tag and reader collision problems create a trade-off between the number of deployed tags (the size of the dataset) and the probability of undetected individuals (the completeness of the dataset). This leads to challenges in inferring missing detections to correct the locations and movement patterns of individuals. Like in other automated data processes, such data imperfections need to be considered and if possible repaired with suitable algorithms.

Here, we demonstrate that non-explicit detection data from fixed observation points contain enough information to infer individual behaviour. Taking advantage of the recent developments in deep learning methods, we develop the “RFIDeep” workflow to automatically extract breeding status and outcome from detection data acquired by RFID antennas using convolutional neural networks. We illustrate how deep learning methods detect biological features in RFID data with very high classification accuracy, and demonstrate the use of a visualisation method not yet commonly implemented in ecology.

RFIDeep was initially developed for an “archetypal” real-life dataset with a ca. 25 years-long RFID detection time series collected on known-age/history king penguins (*Aptenodytes patagonicus*) (Gendner *et al*., 2005). Unlike flipper bands used until then, which are detrimental to the individuals (Gauthier-Clerc *et al*., 2004; Dugger *et al*., 2006), the recording of every transit between the colony and the sea of RFID-tagged birds, throughout their life, allowed a more accurate and unbiased description of the reproductive patterns of the species (Descamps *et al*., 2002), and of the population’s demographic parameters (Le Bohec *et al*., 2008). In these previous studies, all RFID detections were manually analysed by human experts and none of them used the entire dataset of RFID-tagged penguins. Since breeding king penguins exhibit highly stereotyped movement patterns (Descamps *et al*., 2002), they were good candidates for artificial intelligence classification. Based on direct field observations and molecular sexing data, we trained several CNN to infer RFID-tagged penguins’ sex, breeding status and outcome (Breeding vs. Non-Breeding; Success vs. Failure), and breeding dates. We developed RFID-specific data augmentation steps to account for biological variance and data acquisition imperfections. We trained our classification process with field observation data and tested it manually with annotated data to compare the performance of automatic classification with the human experts. We provide all source codes used in RFIDeep workflow that could be applicable for any study using RFID data acquisition and that could inspire ecologists to develop their deep learning process. Finally, a software named Sphenotron, developed to represent movements and locations (in or outside the breeding site) based on RFID detections, is provided with a sample dataset as an example of an RFID data visualisation method.

## 2 Materials and methods

### 2.1 Overall structure of RFIDeep workflow

Figure 1 summarises the steps needed to classify RFID data within a deep learning framework and provides a comprehensive view of the use of the RFIDeep workflow.

**Figure 1:**
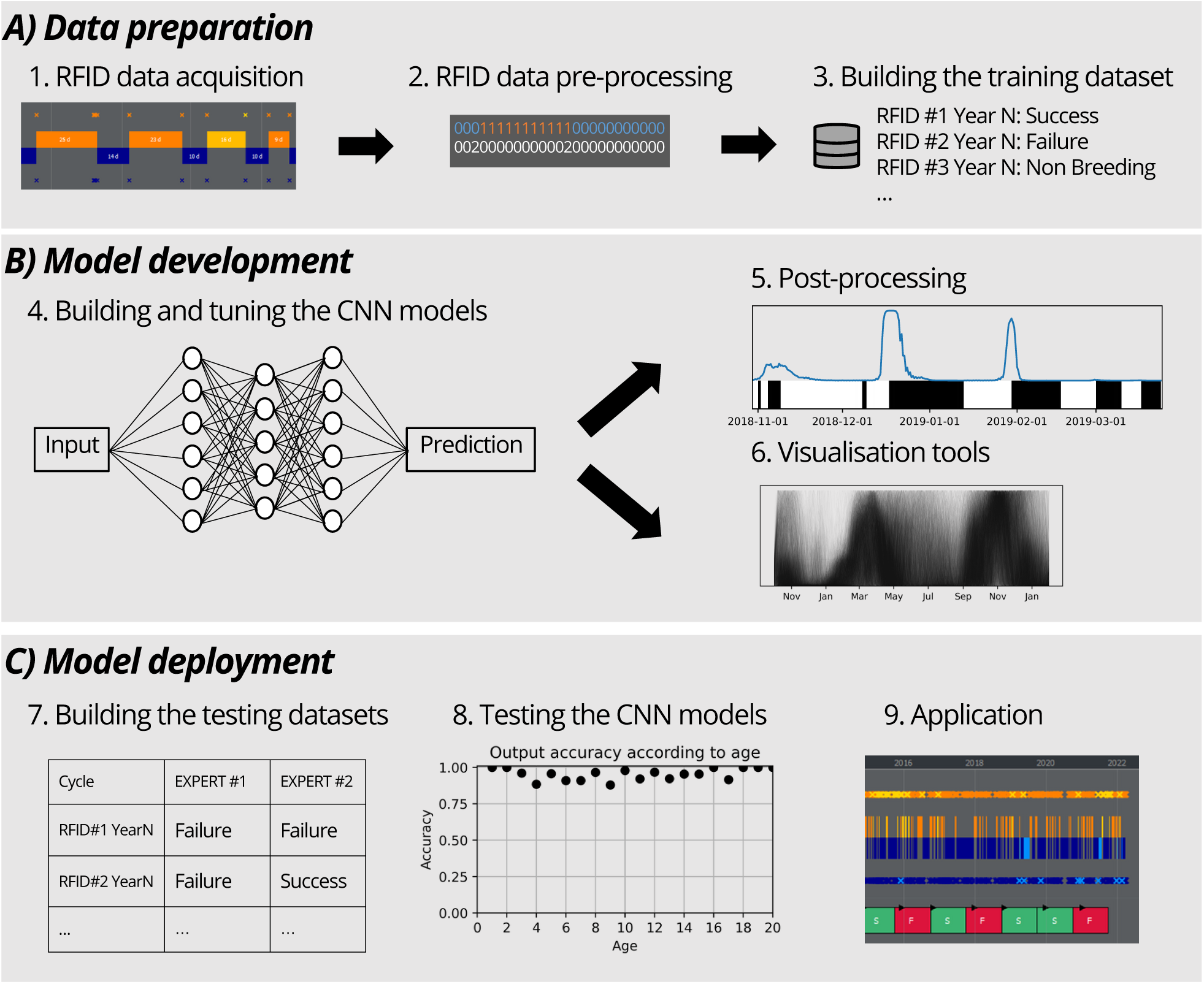
Overall structure of the RFIDeep workflow classifying RFID data with deep learning. The workflow is divided into three phases: data preparation, model development, and model deployment. A) **Data preparation.** 1) RFID data acquisition: many individuals are equipped with RFID tags and antenna systems are installed at key locations to register the detections. A software called Sphenotron (Supplement A) has been developed to represent detections and transitions (in or out of a specific location, e.g., a colony or a nest) of RFID-tagged individuals. A coloring scheme is available to picture the ins (orange) and outs (blue) of tagged individuals. 2) RFID data pre-processing: a correction of missing detections is first applied. RFID data are then formatted analyses (e.g., in or out of a specific location encoded as 1 and 0, and number of detections per time period) to have a unique and readable format for deep learning or for other analyses. 3) Building the training dataset: direct observations of RFID-tagged individuals are used to build a ground truth dataset of labelled vectors giving the true classification. B) **Model development.** 4) Building and tuning the CNN models: the architecture of deep learning models and hyperparameters are tuned with the training dataset. Data augmentation is implemented to cover more biological and technical variance. An individual network is built for each classification problem (e.g., breeding status, sex). 5) Post-processing: classification networks are derived to extract other biological information requiring a post-processing step such as location of stereotyped patterns in RFID data (e.g., determination of the breeding dates with a probability curve (in blue) over presence/absence pattern in black and white, respectively). 6) Visualisation tools: models are validated and interpreted with visualisation tools (e.g., with black curves representing the focus of the model during the breeding season). C) **Model deployment.** 7) Building the testing datasets: a testing step is used to remove biases induced during parameterisation with comparison of model classifications and manual techniques of data processing (i.e., human expert classifications). 8) Testing the CNN models: model tests using expert-labelled datasets assess performance but also ensure that model performances are consistent according to classes and individual characteristics (e.g., age, sex, life stage). 9) Application: classifications are applied to all detection data after pre-processing and formatting (i.e., after correction of missing detections and building of vectors), and results are represented in Sphenotron for each individual (successful breeding cycles in green, failed breeding cycles in red).

### 2.2 Application on a seabird species long-term monitored by RFID

#### 2.2.1 RFID data acquisition

Here, we used data collected from the colony of king penguins (*Aptenodytes patagonicus*) named ‘La Grande Manchotière’ and located at Possession Island, Crozet Archipelago (46°25S, 51°45E). A sub-area of the colony of ca. 10,000 breeding pairs has been electronically monitored since 1998 with RFID technology. As of 2022, four pathways between the sea and the colony (the only ways in or out of the colony) are equipped with permanent automatic identification systems (the detailed information of the field site and systems are described in Gendner *et al*. (2005)). In short, these automatic systems are composed of paired antennas that record the direction of each commuting bird that has been implanted subcutaneously with an RFID tag. Patterns of presence and absence of ca. 15,000 RFID-tagged birds throughout their breeding seasons and life have thus been recorded since 1998. This has generated a large (and increasing) quantity of data, with, for instance, 7 million individual detections as of 2022. To manage, visualise and use information in the field (e.g., select specific groups of birds of known age or history), we developed a python software, called Sphenotron, that displays the location (in or out of the colony) of the individuals during their life, based on the latest known location transition (entrance or exit) for each bird (see Supplement A). Thanks to well-known king penguin’s stereotyped presence/absence patterns at the breeding site (see Figure S7), we can classify the breeding status of any RFID-tagged individual (Non-Breeding, Failed Breeding, Successful Breeding). Start of a breeding cycle (breeding date) can also be determined, it is defined as the beginning of the pattern characteristic of the courtship and incubation period, i.e., the first long sojourn at the colony following the annual moult and exceeding 10 days (Descamps *et al*., 2002). Additionally, the sex of an individual can also be derived from presence/absence patterns at the colony. An automatic sex determination has great potential application for many species where sex determination is challenging (i.e., monomorphic species like king penguins; (Kriesell *et al*., 2018)).

#### 2.2.2 RFID data pre-processing

##### Input data

To prepare the detection data in an appropriate format, we chose to represent absence and presence time-series for each breeding cycle with two vectors providing the location of the individual at the end of 12-hour periods (states 0 and 1) and the number of detections occurring during the 12 hours (Figure S8). For one individual and one given year n, we built vectors encompassing the breeding cycle. For the King penguin, vectors start October 1st of the year Y and end January 31st of the year Y+2 to cover the entire >1-year breeding cycle of the species (Figure S7). We obtained two vectors of 974 elements for each individual and each year. This step can be tailored to match other RFID acquisition systems and species, for instance by dropping the first vector when no location (‘in’ or ‘out’) is defined but only simple detection are recorded, e.g., at a feeding site.

##### Missing detection correction

To tackle missing detections that can occur when individuals exit or enter their breeding site, we developed an algorithm to repair simple missing detections (i.e., detections on only one antenna of a pair, resulting in uncertainty in the individual’s walking direction). These corrections are usually trivial: for example, when an individual is detected only on the inside antenna, followed later by an entrance (i.e., outside-inside transition), an outside detection is inferred to restore a valid pattern in detections corresponding to the missed exit from the colony. We simply built the algorithm to detect all unrealistic successions of detections and to add the corresponding missing detections in all possible cases (See Supplement B).

#### 2.2.3 Building the training dataset

To build a training/ground truth dataset, we visually monitored 295 RFID-tagged individuals over 9 years (2011-2019), assessing their breeding status and behaviour directly through field observations. Birds were monitored from the beginning of the breeding season (November-January), thereby we were able to detect early breeding failures that may have been difficult to distinguish from non-breeding behaviour using RFID detections alone. Breeding outcomes (S: Success; F: Failure) from these study birds was determined according to the survival of their chicks until they fledged. The sex of individuals was determined with the observation of their first period in the colony, as females leave the breeding site right after egg laying, while males care for the egg (Barrat, 1976). A ground truth database with breeding status, timing of breeding, and sexing for 463 breeding cycles was then compiled over the years.

#### 2.2.4 Building and tuning the CNN models

Several models were built to describe breeding activities from regular movement patterns with a classification workflow (see Figure S9):

1. two models to determine if an individual in a given year was a breeder (Breeding vs. Non-Breeding) and if the breeding cycle was successful (Success vs. Failure),
2. a model to distinguish the sex of an individual through sex classification of only breeding cycles and a prediction compiling all the sexes identified over the lifetime breeding seasons,
3. a model to determine the most likely breeding date of males and females separately, through post-processing of a CNN model.

CNN architecture of these models was chosen using a classical simple architecture (see LeCun *et al*. (2015) and through trial and error (see details in S10). Each model was trained on a training set of 80% of the dataset, and the remaining 20% was used as a validation set to measure model performances and avoid overfitting (shown by low validation accuracy and high training accuracy), as suggested by Christin *et al*. (2019). Multiple training of the models with random splitting of the training/validation sets was performed to cross-validate the hyperparameters. Once the final hyperparameters were chosen, validation accuracy with the 20% validation set was recorded, and the final models were trained with 100% of the training datasets. When the models were applied to detection vectors to generate the classifications, the most probable class was chosen for the classification.

To extend the generalisation capacities of our models, we used a data augmentation process during the training of the models (LeCun *et al*., 1998). We used two types of augmentation: the first one consisted in shifting the breeding cycles by a random number of days, as usually done with imagery data to make the models translational invariant. At each iteration of the training, we shifted each training vector by a zero padding at the end or at the beginning of the vector, while trimming the same number of elements on the opposite side. We used a random offset between -30 days and 30 days to cover large biological variability in the phenology of our species (but this can be easily changed in the code, and this step can be turned off). The second augmentation process consisted in simulating missing RFID detections. In the actual dataset, the most frequent problem is the loss of a single detection, which is solved by our correction algorithm. Therefore, we chose to remove 10% of the detections at each iteration, before applying our correction algorithm, allowing a complete recovery of the original detections for at least 50% of individuals (see Supplement C) and leaving uncorrected detections and erroneous locations to improve training generality. Models for determining the breeding status were trained with and without these data augmentation processes to assess the benefits of this step.

#### 2.2.5 Post-processing

### Sex determination

With RFID detections, a human expert can only distinguish males and females based on a few features at the beginning of the breeding cycles, therefore we thought it safer to assume that prediction over a single breeding season would be less reliable than prediction over the whole lifetime. We then averaged the classification probabilities for each sex, for each identified breeding attempt and we obtained the most probable sex over the lifetime of the individuals. We also registered the sex classification for each breeding cycle to measure the benefit of this pooling in classification performance. This step is specific to King penguins and can be skipped for species where sex is easily identified.

### Breeding date

We used CNN models to determine the breeding date by scanning all possible breeding cycles in a year and determining the most probable one (See Supplement D for details on the method). We obtained a certainty curve along the year, with the maximum corresponding to the most probable breeding date (as illustrated in Figure 2 with two true breeding cycles). In our King penguin study case, we trained two different models for males and females separately to account for the difference in patterns at the beginning of their breeding cycles.

**Figure 2:**
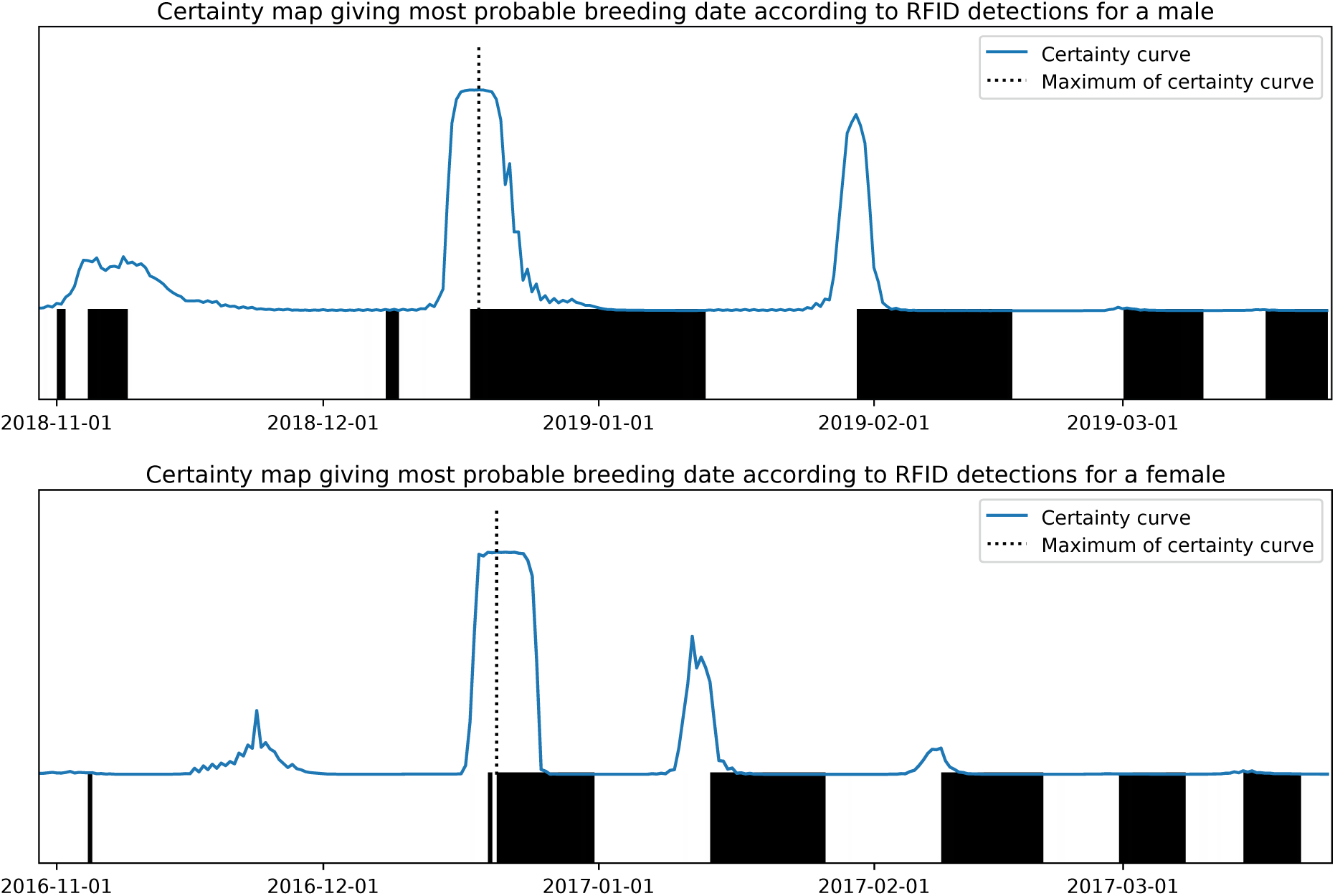
Examples of certainty maps produced by the scanning algorithm to detect the beginning of a stereotyped pattern. Here, the most probable breeding date of a male and a female was determined. The blue curve represents the probability (between 0 and 1) that the breeding cycle starts on a relevant date. The black and white bars in the lower part of the figures represent the location of the RFID-tagged individual (inside and outside, respectively). The most probable breeding date corresponds to the maximum of the blue curve (dashed line).

#### 2.2.6 Visualisation tools

We used visual explanation techniques to show parts of the input data that are identified by the convolutional layers and used to perform the classification. We leveraged techniques recently developed to produce heat maps on images classified by a 2-D CNN algorithm to show which pixels contribute most to the classification (saliency maps (Simonyan *et al*., 2013; Zeiler & Fergus, 2014)) and class activation mapping (Zhou *et al*., 2016)). We produced this type of visualisation on breeding cycles with the GRADient-weighted Class Activation Mapping (Grad-CAM) algorithm (Selvaraju *et al*., 2017) that was directly applicable to the 1-D CNN layers. In short, the Grad-CAM uses the gradients of the final convolutional layer to produce a coarse localization map from an input image (or vector) by searching for pixels whose intensity should be increased to increase the probability of a given class. We ran this algorithm on all breeding cycles in our dataset and obtained a graph of importance value for each element of the vector (each 12-hour period in our example) for a particular class of interest. We computed these activation plots and compared to the raw input detection data for 1) the Breeding vs. Non-Breeding model, restricting the dataset to breeding cycles classified as Breeding in order to identify the features used by the algorithm to detect a breeding cycle, and 2) the Success vs. Failure model, including only breeding cycles classified as successful in order to identify the regions of the breeding cycle that were indicative of a success.

#### 2.2.7 Testing the CNN models

To compare the overall classification performance, we used a global accuracy metric of the different models given by 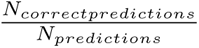 (Powers, 2020). Since our ground truth datasets were well balanced across classes (168 Non-Breeding; 131 Failure; 164 Success), the global accuracy metric did not approach its limits through class unbalance, and it provided a simple and effective metric of overall classification performance. To provide a measure of classification accuracy for all possible classification thresholds, we also used the AUC-ROC score (Area Under the Receiver Operating Characteristic Curve) (Fawcett, 2006). To assess the accuracy of breeding date determination, we used a threshold of 5 days between the true breeding date and the predicted date to define whether or not a breeding date was correctly predicted (see Supplement E). To quantify an unbiased estimate of model performance, we used a testing dataset encompassing 917 breeding cycles of penguin individuals that were never used in model training (Kuhn *et al*., 2013). These breeding cycles were blind-labelled, i.e., breeding status and breeding date were not determined through field observations, but by human experts who examined the RFID detections of individuals using our custom-designed Sphenotron software (see Supplement A). Human experts, with a strong knowledge and experience of the species in the field, were trained using the ground truth dataset, blindly examining detection data to infer breeding cycles, and cross-checking previously assigned breeding cycles. Two human experts were chosen to label the same dataset, and we tested our models with both classifications. We also computed the global accuracy metric between the datasets labelled by the two human experts to assess human variability in classification. The performance of the lifetime sexing method was compared to a molecular sexing dataset of 6,196 birds (molecular sexing method adapted from Griffiths *et al*. (1998) showing 100% accuracy in typical cases (Purwaningrum *et al*., 2019)). Because sex was estimated with a variable number of breeding cycles between individuals (we used all available breeding cycles for each bird), we also tested whether the accuracy of pooled sexing increased when including additional breeding cycles. Finally, we computed the accuracy of the models for each age class and for males and females separately to test whether the performance of our models was consistent over the whole dataset.

## 3 Results

### 3.1 Model training

We chose 200 epochs (i.e., training iterations) for training of each model, which yielded the best results for validating model accuracy while avoiding overfitting. Each model took approximately 1 hour to train using a laptop computer with a CPU Intel Core i7-10750H (2.60GHz) and a GPU Nvidia GeForce GTX 1660 Ti, a non-prohibitive technology as of 2023. The performance of models, according to the validation datasets used to select the CNN architecture and hyperparameters, reached near perfection for the three models, i.e., Breeding vs. Non-Breeding, Success vs. Failure, and Male vs. Female, with global accuracy of 99.1%, 99.7%, and 100%, respectively. As expected, the three models without a data augmentation step achieved lower performances with global accuracy of 94.6%, 91.5%, and 96.6% for Breeding vs. Non-Breeding, Success vs. Failure, and Male vs. Female, respectively.

### 3.2 Model visualisation

Activation maps 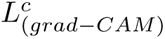 (Figure 3) showed that for the vectors classified in the Breeding class (i.e., individuals that attempt to breed) (Figures 3A and 3C), the model focused on the beginning of breeding, when long periods in the colony occur for breeders but not for non-breeding birds that do not have long fasting periods on the breeding site. For the Success class (Figures 3B and 3D), the model focused the pre-winter period and the post-winter feeding period. As expected, these are the parts of the breeding cycle that can be missing if the breeding fails during incubation, brooding, or even during the winter fasting period. Unsurprisingly, the visualisation maps relied on the same regions that human experts reported using as criteria for determining whether individuals actually attempted to breed and succeeded in breeding.

**Figure 3:**
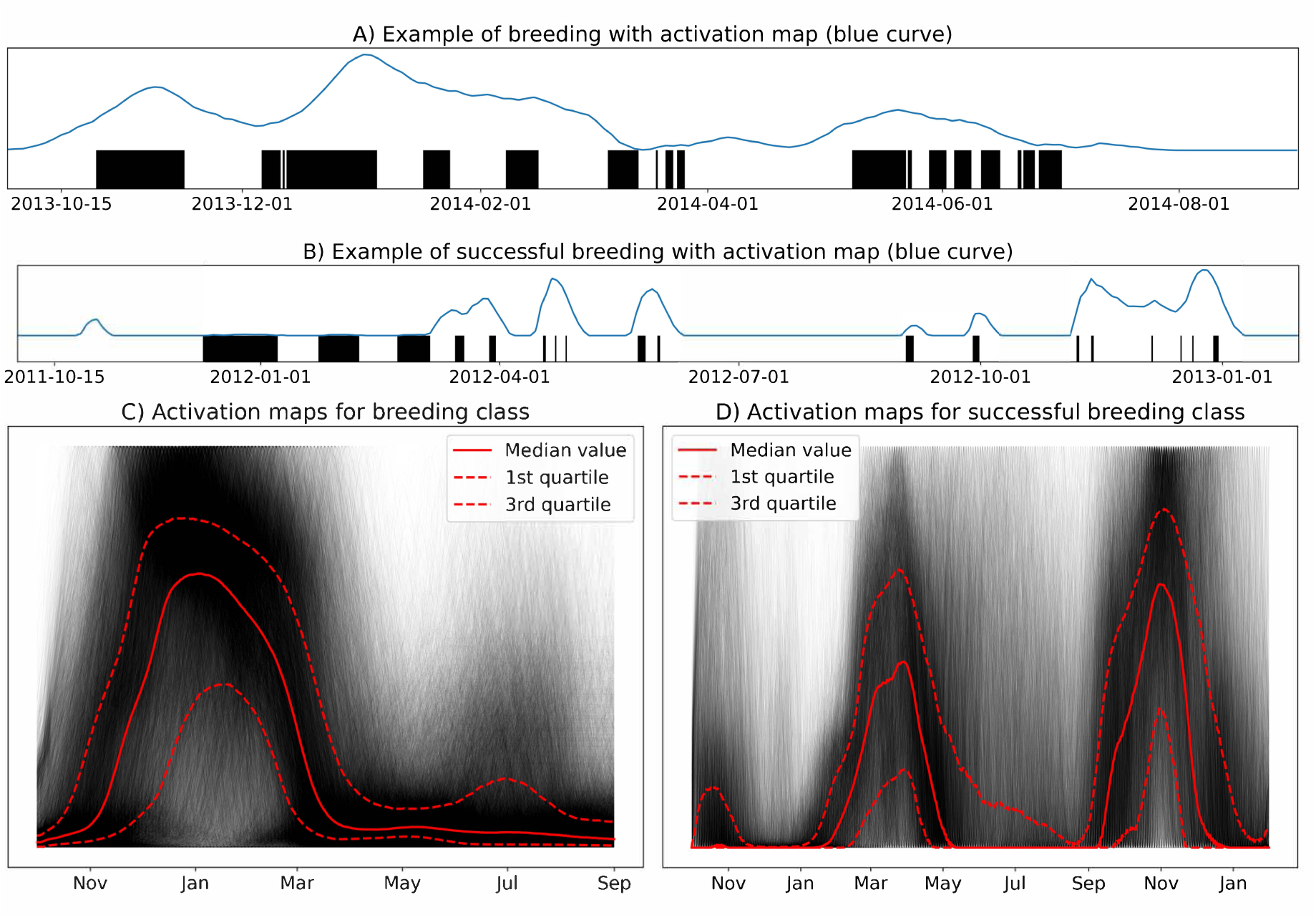
Activation map (blue curve) showing regions of the cycle used by the CNN procedure to produce the classifications and simplified presence/absence pattern (black: inside; white: outside) for two true breeding cycles (A and B), and median maps (C and D) illustrating all maps with the median curve, the first quartile, and the third quartile. A and C correspond to the Breeding class (all breeding cycles classified as Breeding) and B and D to the Success class (all breeding cycles classified as Success), respectively.

### 3.3 Model deployment

The trained models were used to predict the breeding status and dates of all RFID-tagged individuals since 1998 (i.e., 85,524 breeding cycles from 14,795 different individuals). On the laptop computer used here, prediction (from raw RFID data to classification) of breeding status and sex of all birds required 140 seconds, but it took 1.1 hours for the determination of the breeding date due to the number of predictions needed (320 for each breeding cycle). In comparison, it took a human about 1 minute to make the same decision as RFIDeep for one bird and one breeding cycle, which would correspond to 1320 hours or 165 workdays (8h per day) to classify all breeding cycles. Tests of model’s classifications against two human expert classification datasets resulted in high global accuracy metric (Table S1, i.e., 0.963 for Breeding vs. Non-Breeding and 0.967 for Success vs. Failure for Prediction vs. Dataset 1). Similarities between the expert-labelled datasets were globally equivalent to the accuracy of our CNN models, indicating the high performance of the automatic classification procedure. The AUC scores of Breeding vs. Non-Breeding and Success vs. Failure computed with the human expert classification were even higher (e.g., *AUC_Bvs.NB_* = 0.993 and *AUC_Svs.F_* = 0.992 for Prediction vs. Dataset 1, see Figure S11). As expected, models with data augmentation consistently performed better than models without any transformation of the input data. The lifetime classification sexing procedure yielded an 88.6% accuracy compared to the molecular data (*AUC_sex_* = 0.930 see Figure S12). Before pooling lifetime sex probabilities, the global accuracy of sexing was only 81.7%. As expected, we also found that sex classification accuracy from the pooling of lifetime sex probabilities increased with the number of breeding cycles used to determine the sex of an individual (see Figure S12). According to dataset 1, predictions were slightly better for males than for females for the breeding status (males: 94.5% vs. females: 93.2%) and inversely for the breeding dates (males: 85.2% vs. females: 88.9%). The breeding dates also appeared to be less predictable for young individuals (see Figure S13).

## 4 Discussion

In this study, we developed and tested a complete workflow based on CNN models to automatically infer behavioural and fitness traits from RFID-tagged animal detection data. Based on a train-test split approach, we showcased the potential of deep learning to adequately replace human expertise in RFID data processing in a much shorter time span. Remarkably, humanlike performance to translate patterns from detection data into meaningful biological parameters was reached with a rather simple CNN architecture and standard desktop computing capacity. To improve results, we used time-shift data augmentation to mimic the variability that could occur due to biological mechanisms (e.g., a shift in breeding dates) and simulated data dropouts to mimic technical constraints (e.g., missing detections). We also developed a post-processing step to extract dates of breeding, and, with a visualisation technique, we identified the regions of the dataset used by the models to classify the breeding cycles. Such a framework can be used beyond our example dataset and help to quickly classify the breeding activities of many individuals, even more so for long-term projects for which pre-processing analysis is very time- and labor-consuming (in our example, we worked on ca. 15,000 individuals over ca. 25 years). Certainly, we do anticipate a wide application for other colonial birds with their many ins and outs of their colonies and with their complex patterns of phenology. For non-colonial species, we anticipate a good performance of our model with likely simpler RFID data (e.g. simple detection at a feeding site), if detection rate is high enough to capture regular patterns. While it is still challenging to successfully transfer pre-trained deep learning from a study case to another (Marcus, 2018), RFIDeep workflow is tailored for any study classifying behaviours based on RFID-tagged animal detections. RFIDeep was successfully tested and used on another species, the Adélie penguin (*Pygoscelis adeliae*), for which breeding is markedly different from our first dataset example, yet monitored with a similar automatic RFID setup. Given that our model performed well for these contrasting datasets (global accuracy of 95.2% for breeding outcome determination and 93.5% for sex, see Supplement F for details), we argue that any RFID-monitored species with stereotyped movements during a given life stage could certainly benefit from the RFIDeep workflow, such as bumblebees (Molet *et al*., 2008), Leach’s Storm-petrels (Zangmeister *et al*., 2009), hummingbirds (Bandivadekar *et al*., 2018), as well as other penguin species (Kerry *et al*., 1993; Chiaradia & Kerry, 1999; Horswill *et al*., 2014). Furthermore, the missing detection correction and data augmentation algorithms implemented in RFIDeep have great potential to tackle uncompleted and/or low-quality datasets, such as those produced by mobile RFID antennas temporarily deployed (Cristofari *et al*., 2018). We are confident that the RFIDeep workflow will help biologists to adopt deep learning applications more easily, either by using the codes directly or by adapting it for their requirements.

Both validation and testing steps showed the high performances of the RFIDeep models, on the one hand, in reference to the ground truth data and, on the other hand, from a human-machine comparison point of view. Even though we developed a software to efficiently display detections and locations (inside or outside the colony) of RFID-tagged individuals during their life (Sphenotron, see Supplement A), the distinction between specific breeding status can be challenging, if not impossible. For example, a confounding situation in our case study happens between non-breeding and failed breeding when the failure occurs very early in the season. By using automatic classification, we standardised the bias among all breeding classifications throughout the years of monitoring through the removal of variability related to potential differences in human expert interpretation. This allowed for remarkably fast extraction of life history parameters of the monitored individuals, necessary to estimate population vital rates (e.g., survival, breeding success) and viability, in addition to other breeding and phenological traits. For example, breeding success inferred with a very good classification accuracy (97.5% of accuracy in the classification of successful vs. non- or failed breeding) can then be used to estimate fecundity rates of the monitored population with high confidence for all monitored years.

Our analysis highlights the benefits of data augmentation to cope with more biological variance than contained in our ground truth data. Data augmentation is commonly used to improve deep learning applications (Taylor & Nitschke, 2018) and sometimes developed in the application of deep learning in ecology (e.g., with image data (Kälin *et al*., 2019) or audio data (Kahl *et al*., 2021)), and has significantly enhanced our classification process. While data augmentation is usually done by adding random noise to the dataset (e.g., in pictures for 2-D CNN classification with image rotations for instance, (Pawara *et al*., 2017)), here we aimed to mimic biological variance and technical limitations of the RFID data acquisition systems. Doing so, we covered a large variance in breeding dates, enabling us to anticipate breeding seasons that could begin earlier or later than those existing in our ground truth data, because of environmental shifts already observed or expected in the coming years/decades (Visser *et al*., 2021). This applies not only for the Sphenisciformes species used in this study, but likely also for other species with a high variance in breeding phenology (De Villemereuil *et al*., 2020). Another interesting aspect of the automatic classification of breeding cycles is the independence among predictions. Indeed, each breeding cycle was analysed without supplement information about the year (e.g., average breeding success, phenological data), the individual (e.g., age, body condition), and previous and future breeding cycles. The breeding classification of lifetime datasets by human experts can induce bias for quantifying the inter-individual and intra-individual heterogeneity in breeding cycles since they are usually not classified independently. However, while there may be an advantage to having independent classifications, the lifetime information may also be beneficial, for instance to better determine the breeding date of the very first breeding seasons that tend to be less predictable for numerous species (see Figure S13). It would also be useful to train CNN models with mixed data (e.g., RFID detections and automated weighting at the detection point) to increase the classification accuracy and/or complexity, and to refine further some of the analyses (e.g., the stage of breeding failure), as it has also been done in other fields (Ahsan *et al*., 2020). With visualisation techniques, we showed which parts of the datasets are mostly used to perform classification. They provided a peek into the deep learning ‘black box’, making the process more transparent for the user, a shortcoming that often prevents its use by ecologists (Borowiec *et al*., 2022). We argue that such a step can help expand the potential of deep learning to describe and analyse ecological big data. In our example, while activation maps are primarily used by CNN for classification, their visualisation allows the detection of the specific breeding activities or features, such as seasonal phenology. In our application on king penguins, the CNN models showed that the presence or absence of pre- and post-winter chick feeding patterns were the most important criteria for predicting breeding outcome. Although these regions can be used for distinguishing between failure and success, it reinforces our interest in using these visualisation techniques not only to understand how our deep learning models work, but also to detect regions of interest in our datasets. It also highlights the use of CNN models that are not frequently found in ecological studies but have great potential, for instance, to detect hidden patterns in large datasets. Moreover, to cope with the recent explosion of big data acquisition due to increasingly sophisticated, miniaturised, autonomous, and powerful data collection instruments (Williams *et al*., 2020), visualisation tools are becoming increasingly important in detecting similar patterns in given classes or differences between similar classes. For instance, identifying parts of the vocalisation essential to distinguish between species or even individuals is key in bioacoustic studies (Stowell *et al*., 2016; Kobayashi *et al*., 2021). Visualisation techniques have also been used to select the most informative variables to infer animal behaviours from multi-sensor data (in green turtles (*Chelonia mydas*) (Jeantet *et al*., 2021)) or to select the most relevant morphological characters to identify species (among midges (Milošević *et al*., 2020) and mosquitoes (Park *et al*., 2020)). By developing tools to help users unleash the vast potential of machine learning in ecology and to increase numerous benefits of RFID technology, we also aim with RFIDeep to foster low-impact monitoring of sensitive species by reducing human presence and intervention in wild habitats (Hughes *et al*., 2021; Rafiq *et al*., 2021; Harrison & Kelly, 2022). We are convinced that combining automatic data collection and real-time data analysis and storage will help secure key ecological information over time necessary to continuously monitor the health of wild populations and their ecosystems.

## Acknowledgements

This study was supported by the Institut Polaire Français Paul-Emile Victor (IPEV) within the framework of the Program 137-ANTAVIA, by the Centre Scientifique de Monaco with additional support from the LIA-647 and RTPI-NUTRESS (CSM/CNRS-UNISTRA), by the Centre National de la Recherche Scientifique (CNRS) through the Programme Zone Atelier de Recherches sur l’Environnement Antarctique et Subantarctique (ZATA), by the Deutsche Forschungsgemeinschaft (DFG) grants ZI1525/3-1 and ZI1525/7-1 in the framework of the priority program “Antarctic research with comparative investigations in Arctic ice areas”, and by Academy of Finland grant #331320. This study was approved by the French ethics committee (last: APAFIS#29338-2020070210516365) and the French Polar Environmental Committee and permits handling animals and access breeding sites were delivered by the “Terres Australes et Antarctiques Françaises” (TAAF). We are deeply grateful to all the wintering and summering members of Program 137, Denis Allemand, Benjamin Friess, Yvon Le Maho, Victor Planas-Bielsa, Claire Saraux, and all the other colleagues and students within the team who have contributed over the last 20 years to the improvement of hardware, software, and databases. We also sincerely thank the IPEV logistics teams for their important and continued support in the field.

## Author’ contributions

G.B., R.C., N.C., M.G.-C., J.-P.G., Y.H., C.-E.S, D.P.Z., C.L.B. conceived the ideas and designed methodology; G.B., R.C., T.B., M.B., N.C., F.A.N.F., M.G.-C., A.H., A.K., N.L., C.-E.S, E.T., B.V., C.L.B. collected the data; G.B., R.C., T.B., M.B., C.C., F.A.N.F., M.G.-C., A.H., C.-E.S, B.V., C.L.B. cured and/or analysed the data; G.B., R.C., A.W., C.C., J.C., A.H., C.-E.S, E.M.W., D.P.Z., wrote codes for RFIDeep and software; R.C., N.C., J.C., M.G.-C., J.-P.G., Y.H., D.P.Z., C.L.B. developed the hardware; Project administration and supervision: C.L.B.; G.B. and C.L.B. led the writing of the manuscript. All authors contributed critically to the drafts and gave final approval for publication.

## Data accessibility

RFIDeep and Sphenotron codes are accessible at the following location: https://github.com/g-bardon/RFIDeep

## Supplement A Sphenotron software

Sphenotron is an open-source software written in Python. The current version of Sphenotron has been developed to interact with MySQL databases composed of several tables compiling all known information on each RFID-tagged individual, such as biometric or phenological data, sampling and recapture events if any, past breeding territories/coordinates, etc., in addition to detections (see Figure S1). Sphenotron was initially developed to interact with databases related to specific species (order Sphenisciformes). Yet, the code can be reused and adjusted to manage and interact with databases of similar or different formats of other species. The complete Sphenotron software and codes, with examples of database, are downloadable at the following link (https://github.com/g-bardon/RFIDeep), and can be fully modified for a wide range of species (e.g., sea or terrestrial birds or mammals, fishes) and monitoring scheme (e.g., with and without mass tracking).

With automatic data pre-processing and analysis, Sphenotron allows the organisation, aggregation, management, and storage of biological time series in near real time (Figure S2). The development, improvement and tests have been implemented to long-term monitored penguin populations since 2002. With the development of this novel interface, we aim to optimise data reuse following the FAIR principles (Wilkinson *et al*., 2016). In addition to automating the time-consuming pre-processing of detection data into biologically meaningful data (e.g., breeding outcome, sexing), the other advantage of this automatisation is to assist scientists in the field by accessing information from RFID-tagged individuals to conduct specific experiments or observations on selected individuals with the desired characteristics.

**Figure S1:**
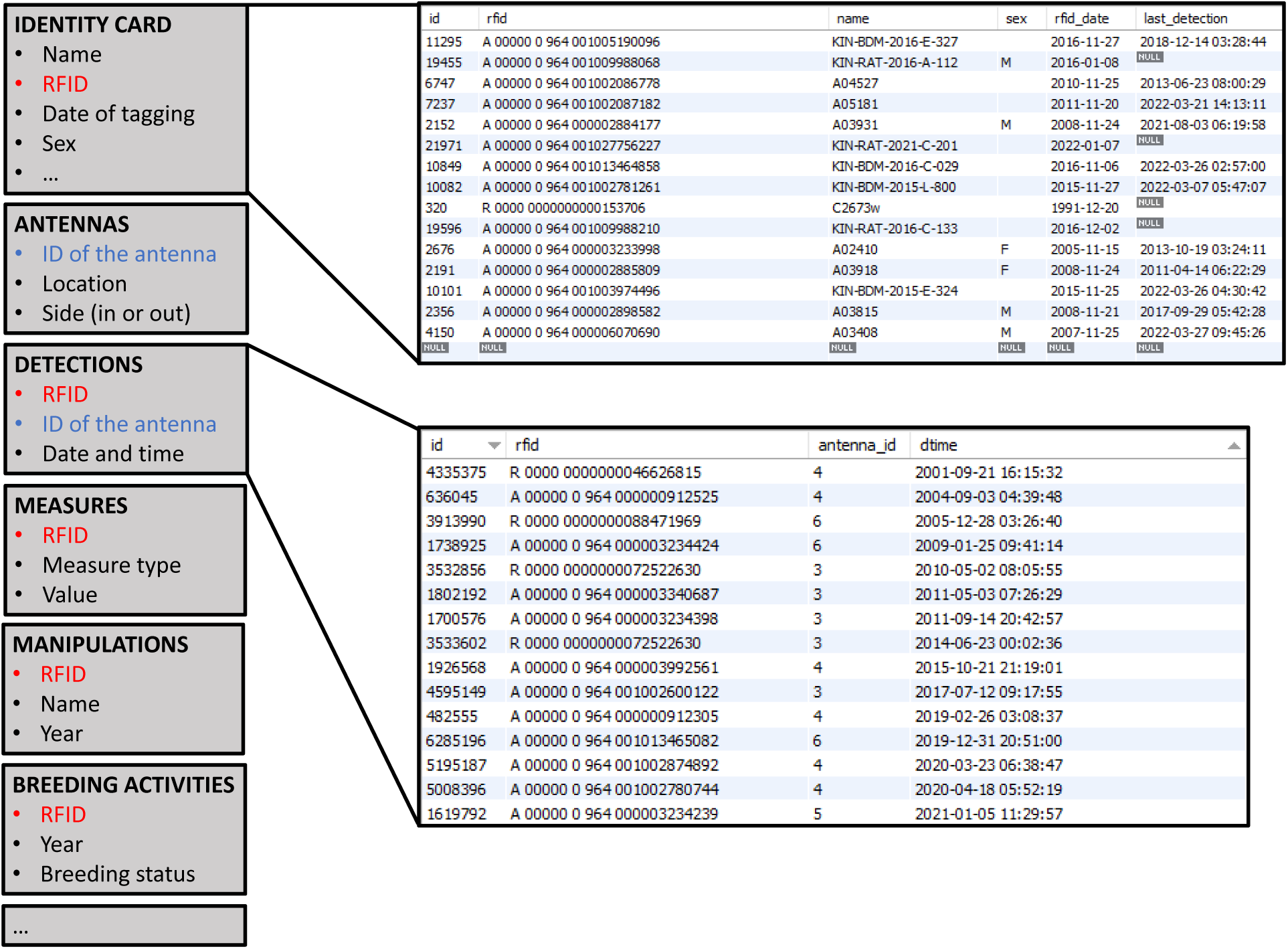
Schematic illustration of the main tables of the database. Each light grey box corresponds to one table. The database is built around the main table (IDENTITY CARD) containing all the key information of the individuals. The other tables refer to this main IDENTITY CARD table with the identity of the individuals given by the RFID numbers. For example, the DETECTIONS table records all RFID detections with the RFID identity of the detected individual and the identity of the antenna, also linked to ANTENNAS table compiling all information about the different antennas. Additional information on the individuals is given in other tables for better clarity and flexibility in data storage. Extract of the IDENTITY CARD and DETECTIONS table are given on the top and bottom right side, respectively. The ‘id’ column stands for the unique identification number assigned to each row of the table. The ‘rfid’ column corresponds to the RFID-tag number (and by extension the identity of the associated individual). The ‘name’ column gives the name of the individuals used in the field for simplicity and clarity. The ‘sex’ column gives the molecular sex if known. The ‘rfid_date’ column gives the date of RFID-tagging. The ‘last_detection’ column gives the date of the last detection on the antennas and is continuously updated. The column ‘antenna_id’ corresponds to the identification number of the antenna. The ‘dtime’ column gives the date and time of the detection.

**Figure S2:**
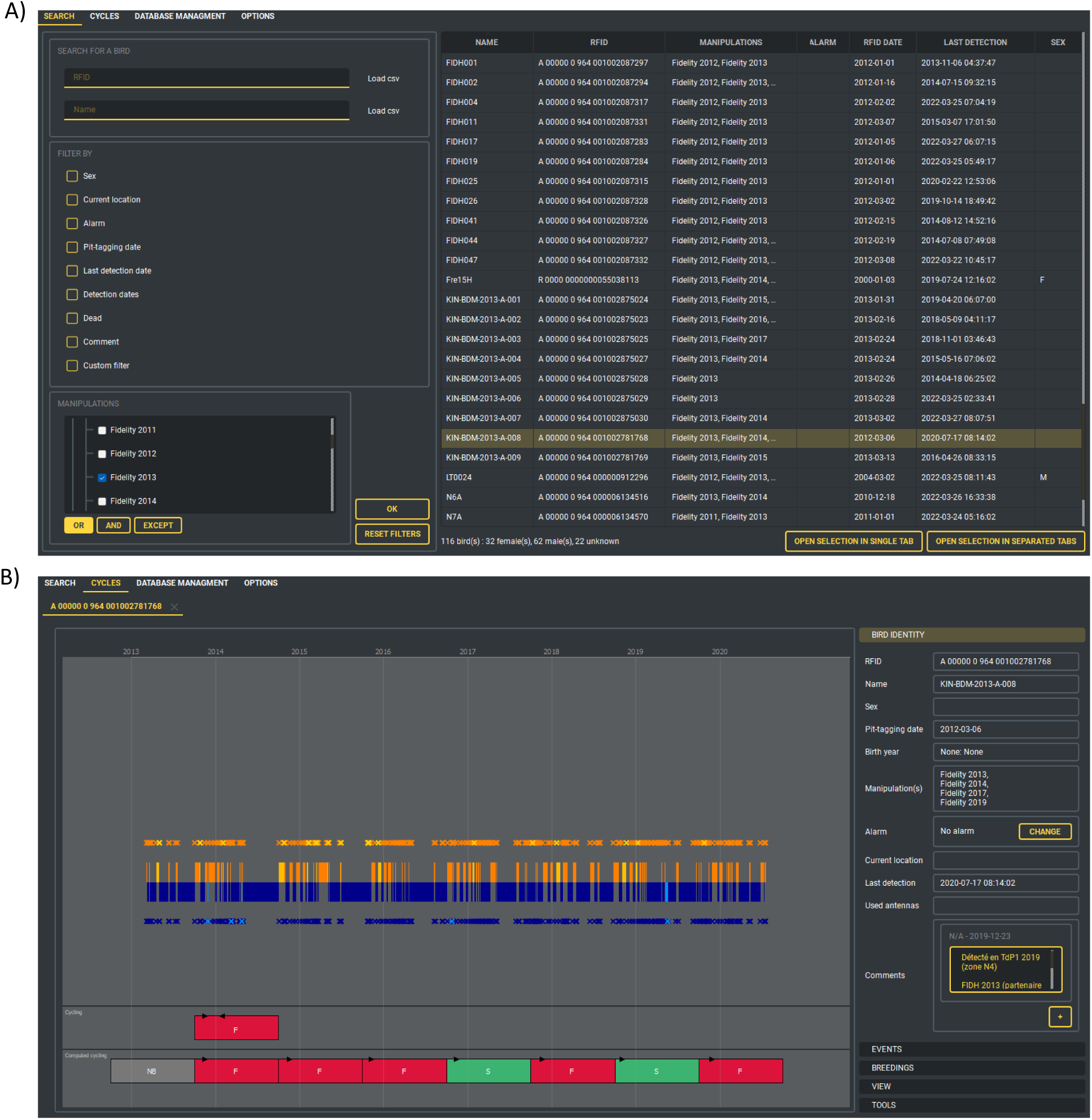
(A) Main window of the Sphenotron showing a search feature to easily find one or more individuals in the database. The left side of the window contains common filters used to search for specific birds according to chosen filters (e.g., filter by Sex, PIT-tagging date, or Manipulations). The right side gives the result of the search in the database and the main information known on the individual. These individuals can be selected, and the detections can be displayed in (B) Individual window showing the presence/absence pattern of a selected individual (here KIN-BDM-2013-A-008) during its electronically monitored lifetime. The right panel displays the individual’s information. The central chart displays the lifetime locations of the individual: inside the colony in orange, and outside the colony in blue. Yellow and light blue correspond respectively to inside corrected location and outside corrected location (with the algorithm of missing detection correction). Each cross corresponds to one detection (with the same colour scheme). The bottom panel displays the breeding cycles analysed by human experts (top) and by RFIDeep function (bottom): Success in green, Failure in red, and Non-Breeding in grey. Black triangles correspond to the Breeding dates.

## Supplement B Missing detection correction algorithm

To build the missing detection correction algorithm, the detections have been converted into short or long transitions (duration between two following detections and the side of both detections), which were coded in 3 bits: the 1st bit gives the side of the first detection (0 for outside and 1 for inside), the 2nd bit gives the duration between the two detections (0 for less than 10 minutes and 1 for more), and the 3rd bit gives the side of the second detection. These encoded transitions have then been converted into numerical and the vectors of all transitions have been built. A correct schema of transition is therefore 1-7-4-2 (corresponding to 001 – 111 – 100 – 010 in 3-bit code) as shown in Figure S3. The algorithm has then been built to detect the incorrect successions of transitions and to correct each possible error by adding one or several detections to recover the 1-7-4-2 successive transitions. Some missing detections remain impossible to correct in this way, e.g., an individual leaving the colony two successive times without entering in between, because no information is known on the date of the entering.

**Figure S3:**
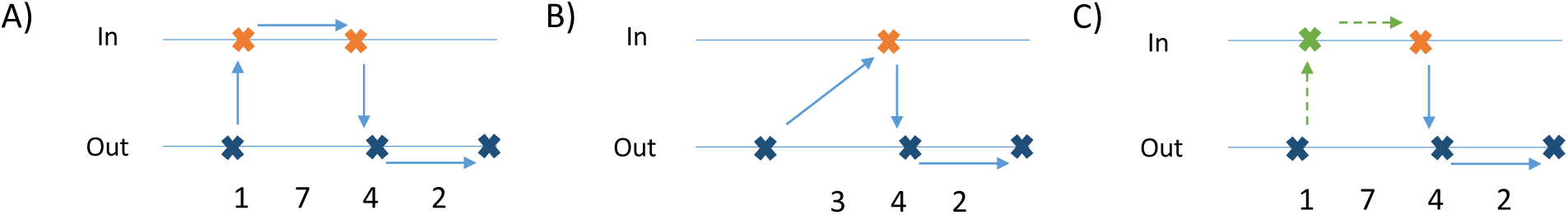
The blue crosses correspond to true detections at the outside antenna and the orange crosses to true detections at the inside antenna. The green cross corresponds to a logically added detection. Arrows represent transitions. A) Correct schema of transition. B) Incorrect schema of transition leading to an impossible long duration between detection time recorded by the outside and inside antennas. C) Schema that gives the correction of B) with the addition of a logical detection (green cross) just after the first one to recover the correct schema.

## Supplement C Missing detection correction performances

The algorithm to solve the missing detections was tested based on the detections from our ground truth dataset to assess its performances according to various degrees of missing detections. The first step was to get a cleaned detection dataset with corrected detections (because we used detections from the field). Thus, detections have been corrected with the algorithm to remove original missing detections and we manually validated the new corrected detections. Then, this cleaned dataset was used to test the algorithm performance by removing a various number of detections and by comparing the cleaned dataset with the newly corrected one (Figure S4).

**Figure S4:**
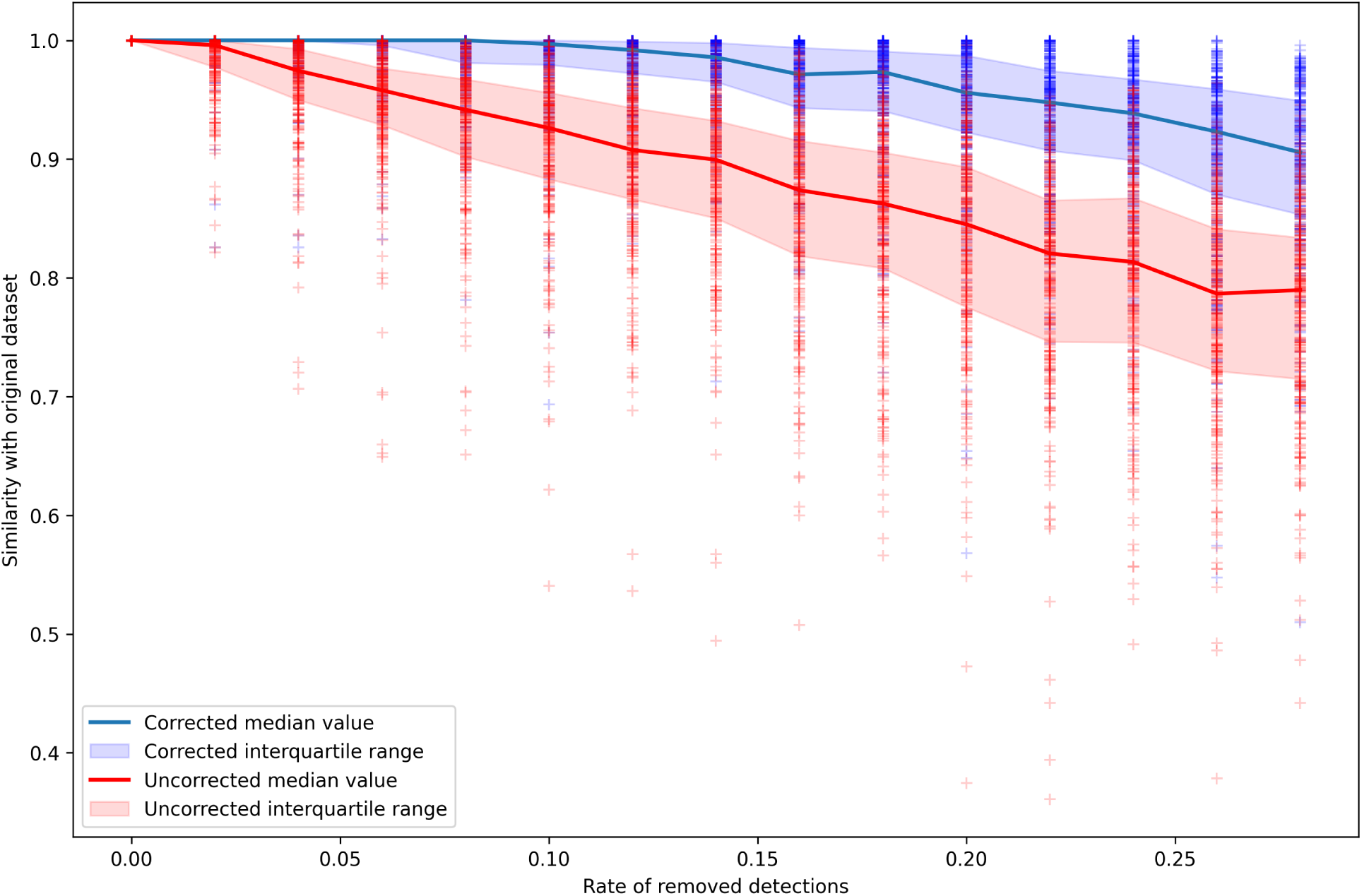
Performance of missing detection correction algorithm with various rates of removed detections. Similarity of location vectors giving the location (inside or outside) of the individuals in 12-hour periods during the breeding season are compared to the cleaned location vector where raw RFID data was corrected by the algorithm. The blue curve gives the median of the similarity of the vectors for the corrected one while the red curve gives the uncorrected vectors similarity.

## Supplement D Breeding date determination

We trained a new CNN model (with the same architecture and hyperparameters as before) to classify whether the detection vector was aligned with the breeding date (positive class) or not (negative class). To build the positive class, a dataset where each vector was aligned to a known breeding date was designed. We used ground truth breeding cycles with a known breeding date and truncated the detection vectors around the breeding date (30 days before and 75 days after). These vectors constituted the positive class of our training dataset with vectors aligned on the breeding date. The dataset was completed with a negative class corresponding to breeding cycles that were not aligned to the breeding date (e.g., starting at an unrealistic date): we draw a random breeding date for each correct breeding cycle and truncated the vectors around this random date, giving us the second half part of our training dataset. This data generation was repeated at each iteration (epoch) of the training to cover the maximum number of unrealistic breeding cycles possible while keeping a 50/50 ratio of positive to negative classes at each iteration. The data augmentation process were tailored to fit with this specific classification: the shift of breeding cycles were turned off because it would make our classification irrelevant. To apply this model and obtain the most probable breeding date for a given breeding cycle, we classified detection vectors that were aligned to each 12-hour period between November 1st to April 1st, and assessed the probability of having a correctly aligned vector with the previously trained model.

## Supplement E Computation of breeding dates accuracy

To compute the accuracy of breeding date determination, we used a score giving the proportion of breeding dates that were correctly determined by taking a threshold of 5 days between the true breeding date and the prediction. Indeed, most predictions are either perfectly accurate or completely false (Figure S5). This threshold was chosen to consider uncertainties in the definition of the breeding date (the beginning of the first long period on the breeding site is not always well clearly recognisable) and because the period of 5 days before and after the predicted date leads to only one possible breeding date given that the first period on the breeding site lasts more than 10 days for our species.

**Figure S5:**
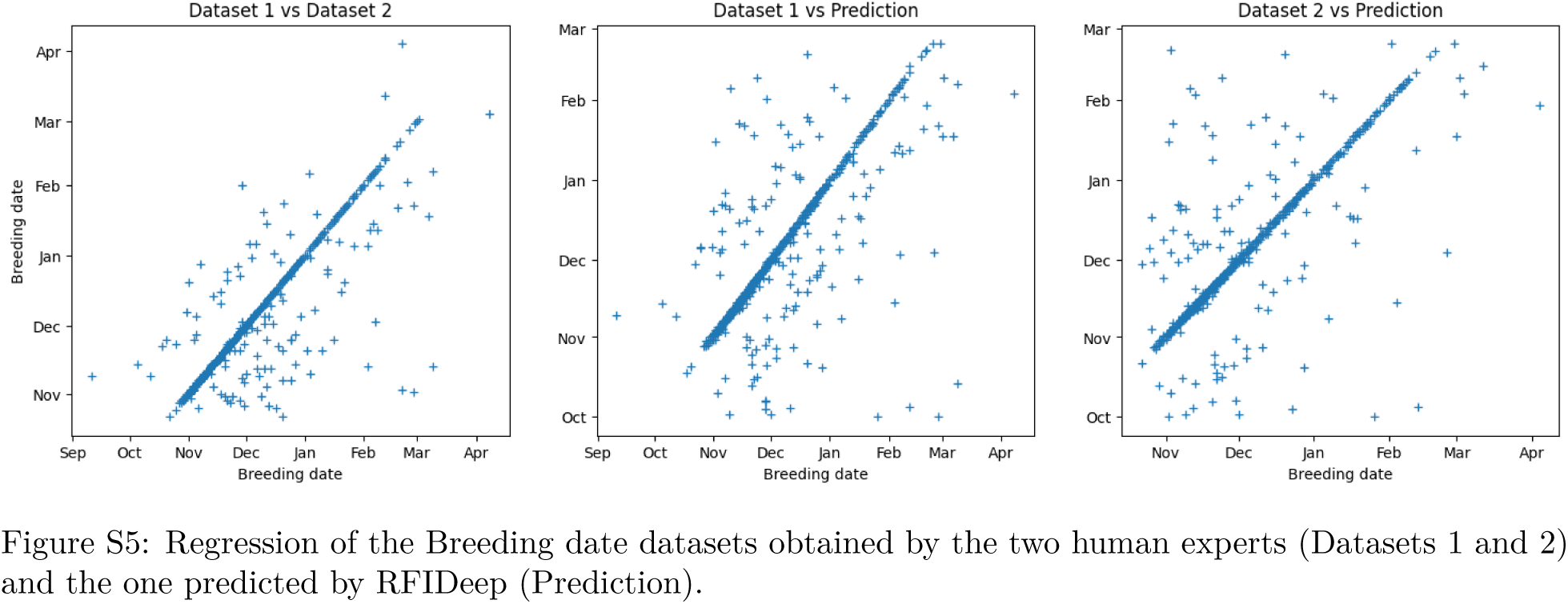
Regression of the Breeding date datasets obtained by the two human experts (Datasets 1 and 2) and the one predicted by RFIDeep (Prediction).

## Supplement F Applying RFIDeep to another species, the Adélie penguin

Similar to king penguins, Adélie penguins (*Pygoscelis adeliae*) routinely perform foraging trips between their breeding colonies and the sea during reproduction. However, their breeding cycle is much more constrained temporally than that of the king penguin. All land-based breeding activities take place during a 6-month window between October and March (Ainley & deLeiris, 2002). As part of the long-term monitoring program 137 of the French polar Institute Paul-Emile Victor (IPEV), an Adélie penguin colony of ca. 300 breeding pairs are electronically monitored with RFID tags off the coast of Adélie Land, Antarctica (ANTAVIA colony, Île des Pétrels, Pointe Géologie archipelago). Every breeding season since 2009-2010, two-access pathways equipped with RFID antennas record the colony attendance patterns of known RFID-tagged individuals. As for king penguins, these patterns are informative on both the breeding outcome (Success or Failure) and the sex of individuals. These biological parameters are not easily determined by direct observations yet are critical for understanding population processes from individual-based data. There is thus a genuine interest in an automated assessment of individual sex and breeding outcome, especially given the potential for comparisons with other locations around Antarctica, where similar electronic monitoring setups exist (Kerry *et al*., 1993; Olmastroni *et al*., 2000; Ballard *et al*., 2001; Lescroël *et al*., 2014; Afanasyev *et al*., 2015). Here, the RFIDeep approach was applied to 3,959 breeding cycles collected between 2009-2010 and 2021-2022 (907 unique individuals). One breeding cycle corresponds to all the detections of an individual in each breeding season (Figure S6). The algorithm was trained using ground truth data from 319 breeding cycles, when breeding outcomes and sex were determined by (labor-intensive) field observations. The resulting model’s accuracy was then tested on a separate dataset of 1,164 breeding cycles, where breeding outcomes and sex were determined by human observation of breeding cycles (e.g., as in Figure S6). In this new application, no hyperparameter in the architecture of the model was changed (i.e., we kept the same number of layers, filters, and kernel), but the length of the input vector (364,2) was modified to fit the shorter breeding season of the Adélie penguins (here it corresponds to 182 days with a 12-hour time-steps, meaning 6 months, encompassing all possible breeding season lengths). The accuracy of RFIDeep reached 95.2% for breeding outcome determination and 93.5% for sex. These values are comparable to those obtained for king penguins. These results further demonstrate the broad applicability and effectiveness of RFIDeep for extracting biologically meaningful parameters from RFID data. This electronic monitoring is also paired with weighbridges, meaning that weights are recorded when an individual crosses the bridge and is detected by the antenna. In future development of the RFIDeep, mass information could be paired with the detection vectors to help the model to determine individual status and sex.

**Figure S6:**
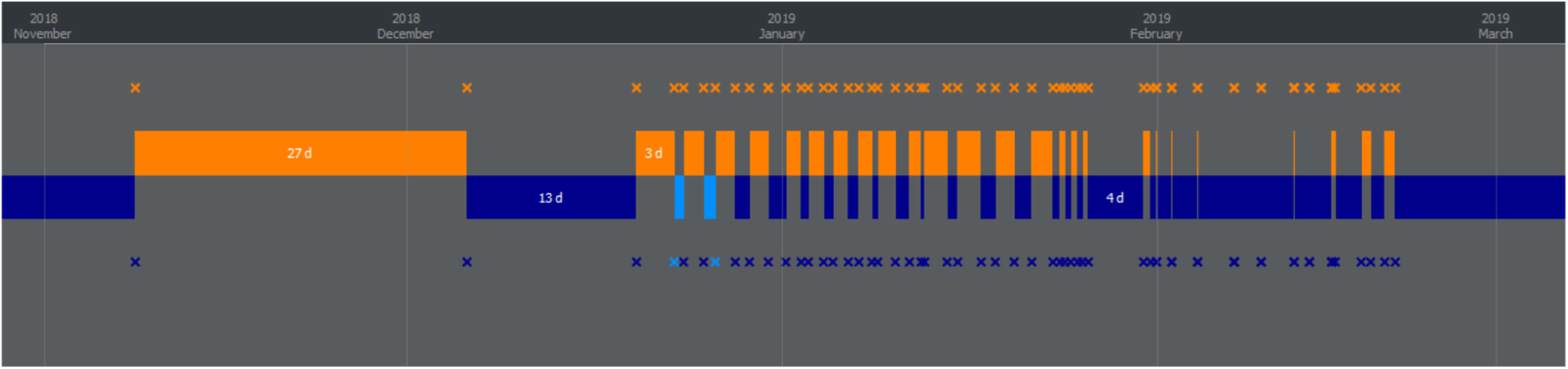
Typical pattern of presence/absence of a successful breeding male Adélie penguin. Each cross corresponds to one detection by an antenna (blue and orange indicating outside and inside, respectively). Light blue indicates corrected detections and locations at sea.

## Figures and Tables

**Figure S7:**
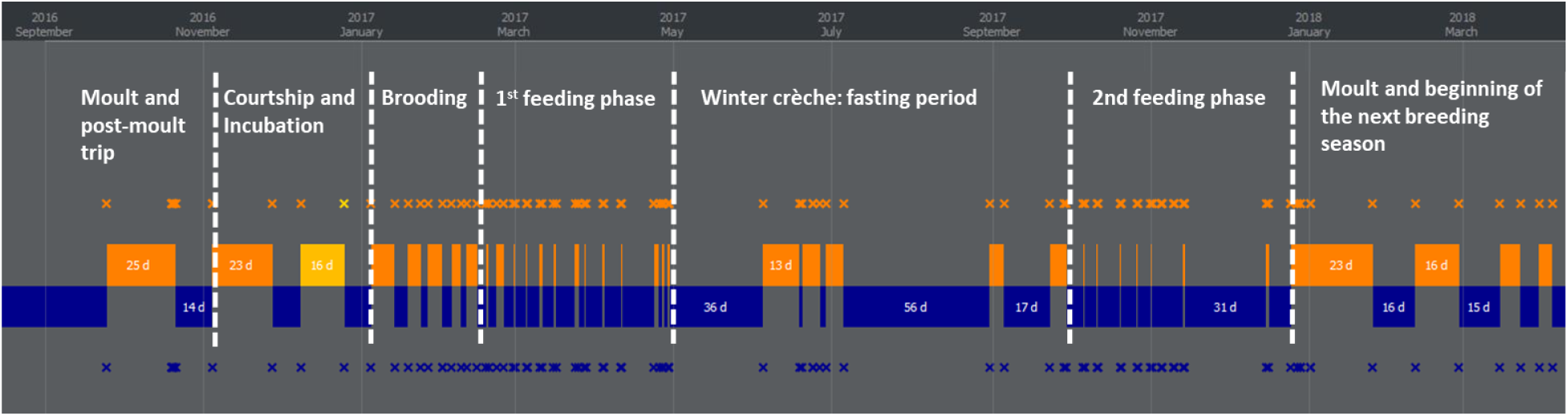
Representation of the presence/absence patterns at the breeding site of a given RFID-tagged individual for one successful breeding cycle. Each cross corresponds to one RFID detection (outside antenna in blue, and inside antenna in orange). The periods outside the colony (in blue) and inside (in orange, or in yellow after the correction of missing detection) are deduced from the sequence of detection. The duration in or out of the specific area is given in days (d). The presence/absence patterns presented here correspond to the annual activities of a male king penguin but can be applied to any individual or any targeted species to understand, for instance, how long an individual stays or leaves a specific study area where RFID antennas are installed. Phases of the breeding cycle, identified from the presence/absence patterns of a bird, are indicated by the white dashed lines. The breeding cycle starts between November and March, and a successful cycle lasts 12 to 14 months (Barrat, 1976). Three status can be listed: (1) Successful breeding with regular presence/absence patterns during the first austral summer and after the austral winter, (2) Failed breeding when at least one major pattern of (1) is lacking, and (3) Non-breeding when no regular presence/absence pattern is identified.

**Figure S8:**
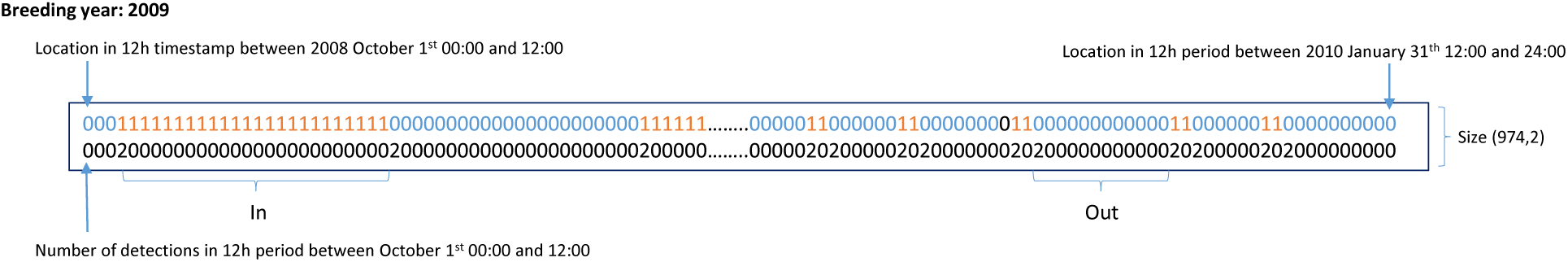
Example of one breeding cycle converted into 2 vectors of length 974. The first row corresponds to the location of the individual at each 12-hour period (0 for outside and 1 for inside). The second row gives the number of detection per 12-hour period.

**Figure S9:**
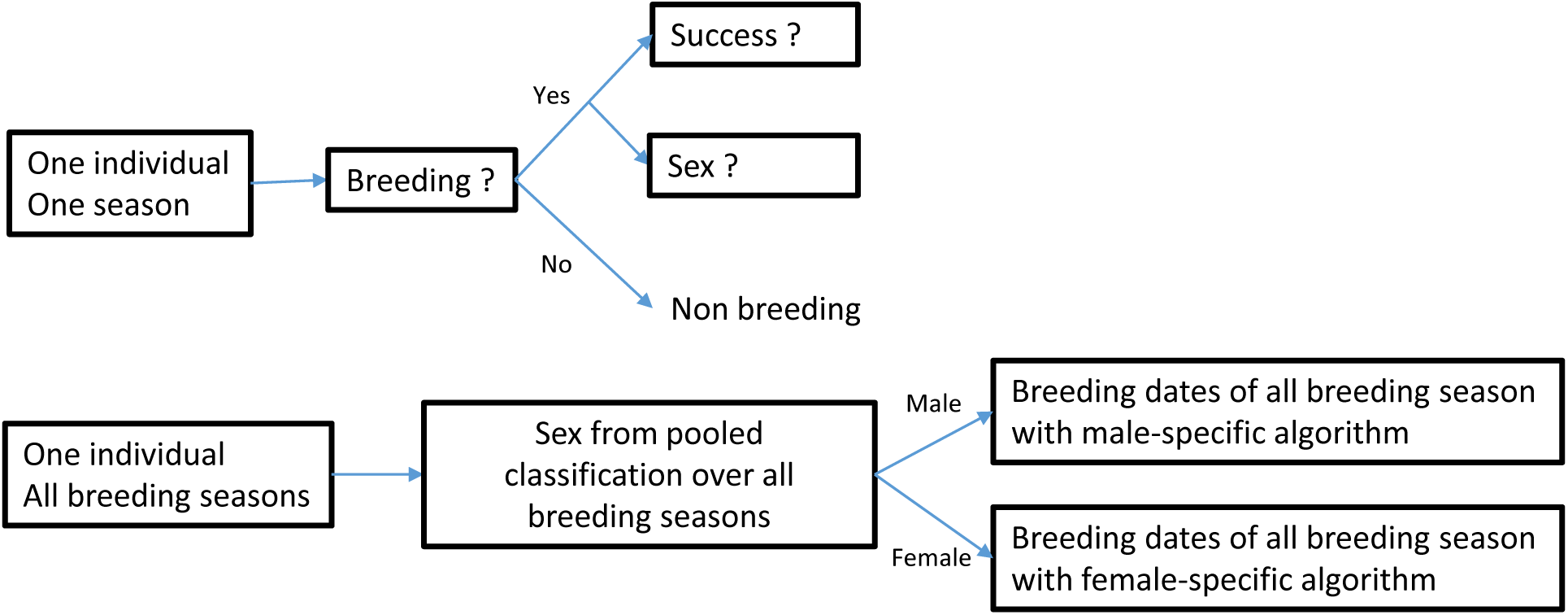
The classification scheme used to determine breeding specifications. The top part of the scheme gives the classification procedure performed on each individual and each season: it leads to classification of the breeding status (Breeding vs. Non-Breeding), the breeding outcome (Success vs. Failure), and the sex (Male vs. Female). The lower part of the scheme corresponds to the determination of sex from the pooled classification of breeding cycles and to the determination of breeding dates of each breeding season according to the sex.

**Figure S10:**
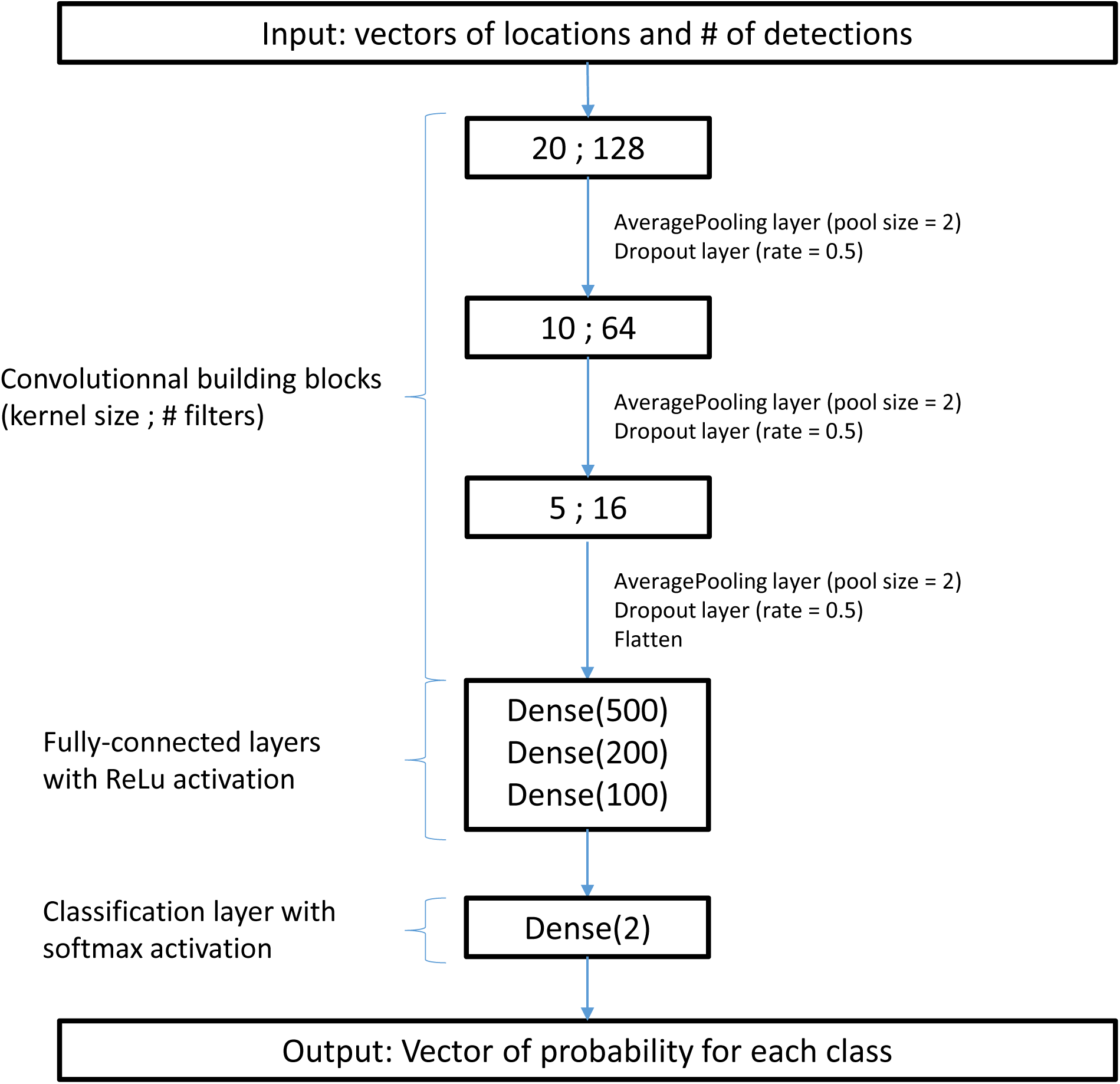
CNN architecture used for the classification model of Success vs. Failure. The CNN architecture consisted of three 1-D convolutional layers (with the size of the kernels and the number of filters shown in the three boxes on the top). Each convolutional layer was interleaved with a dropout layer and a common value of 0.5 as drop rate (to avoid overfitting (Srivastava *et al*., 2014)) and an average pooling layer (with typical pool size of 2) to keep the most essential elements (see LeCun *et al*. (2015) for details on CNN architectures). A flatten function was used at the end of convolution blocks to obtain a single 1-D vector from the previous layers. Three fully connected layers (Dense) followed the building blocks with the Rectified Linear Unit or ReLU (default activation function that applies f(x) = max(0,x) to all neurons, used to improve performance of training) and were included just before the prediction layer to interpret the learned features. The final classification uses the most common softmax activation function with a fully connected layer (Dense) that convert vectors of numbers (the last Dense layer) into a vector of 2 output probabilities. Several CNN models were trained to classify the detection vectors into different classes (Breeding vs. Non-Breeding; Success vs. Failure; Male vs. Female), but the same CNN architecture was used for each classification, meaning that the same layers were used in the same order. Kernel sizes were chosen through trials and errors and multiple training of the model leading to kernel of size 20, 10, and 5 for Success vs. Failure and Male vs. Female models, and 50, 20, and 10 for Breeding vs. Non-Breeding model. The CNN was implemented using the Keras tensorflow framework (Abadi *et al*., 2015) in Python 3.9.7.

**Table S1:**
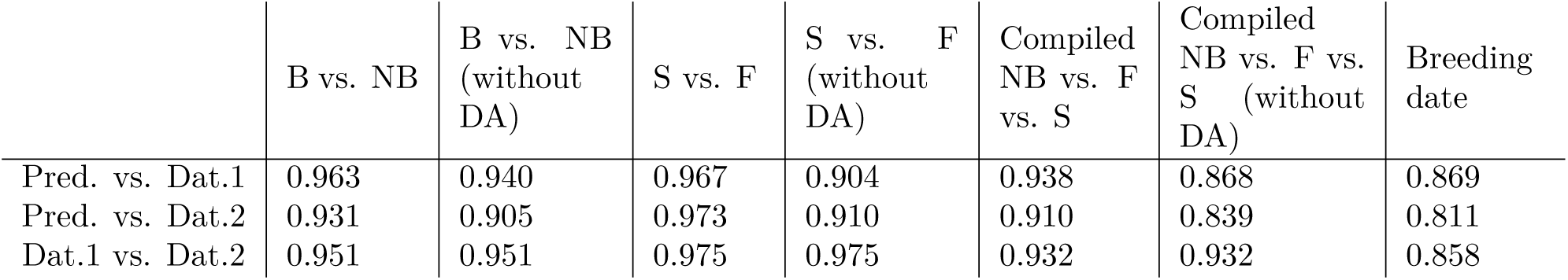
Results of the comparison between the predictions (Pred.) yielded by RFIDeep and two datasets labelled by human experts (Dat.1 and Dat.2). These datasets were well balanced across classes, with about 50% of Failure and 25% of Success and Non-Breeding. For each model (with and without data augmentation ‘DA’ procedure), global accuracy metrics between CNN predictions and the two datasets labelled by human experts are given, as well as the global accuracy between the two datasets. These accuracy metrics are given for the classifications of Breeding vs. Non-Breeding (B vs. NB), Success vs. Failure (S vs. F), compiled Non-Breeding vs. Failure vs. Success (NB vs. F vs. S), and Breeding date.

**Figure S11:**
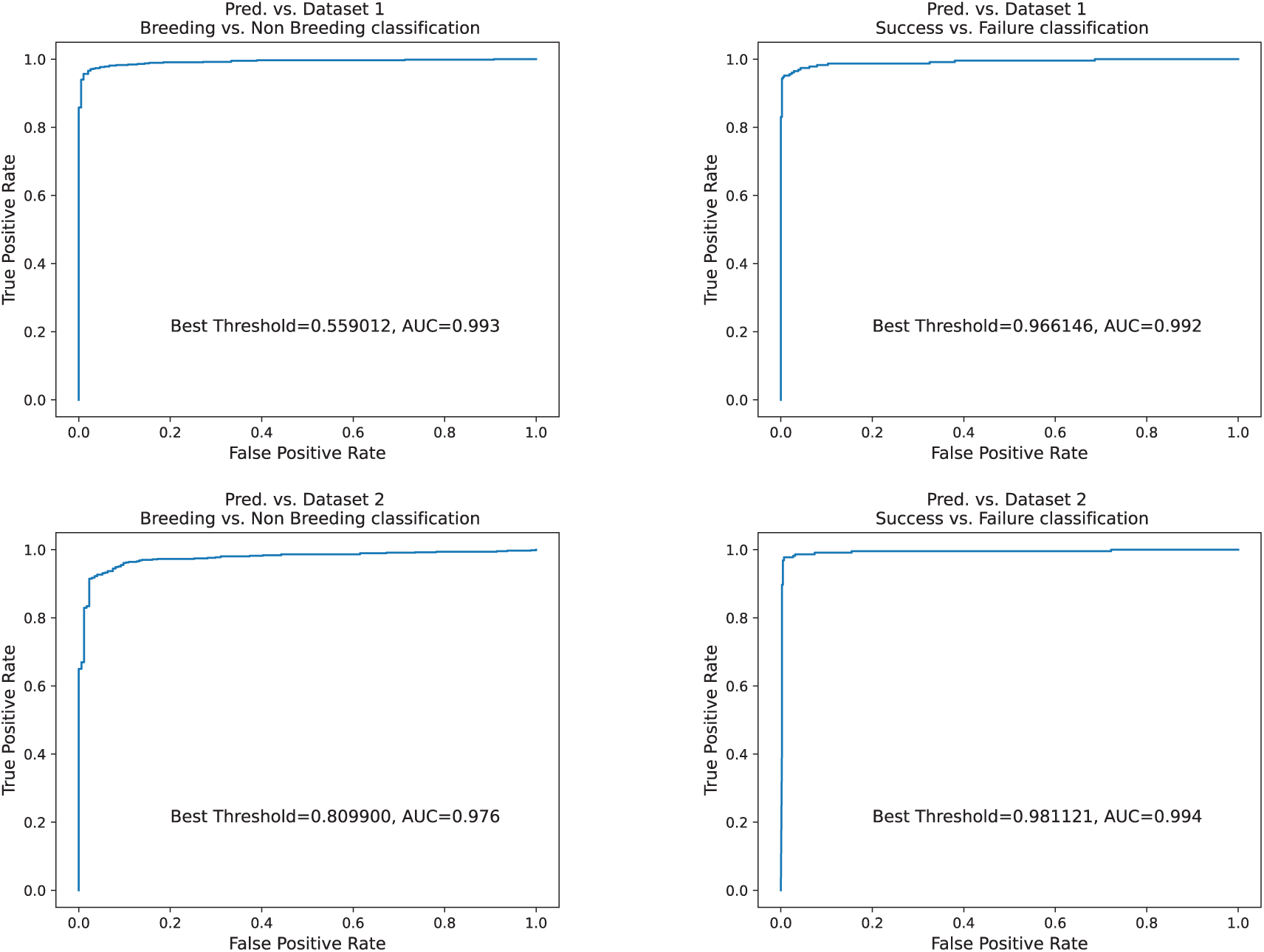
Receiver Operating Characteristic (ROC) curves and associated Area Under the Curve (AUC)-ROC scores, and best classification threshold for the Breeding vs. Non-Breeding model and for the Failure vs. Success model. These metrics were computed using two human expert labelled datasets. High AUC-ROC scores (i.e., > 0.99) reveal high accuracy in comparison to the human performances. The values of the threshold reveal that the models tend to be less restrictive to classify a breeding cycle as a success than the human expert.

**Figure S12:**
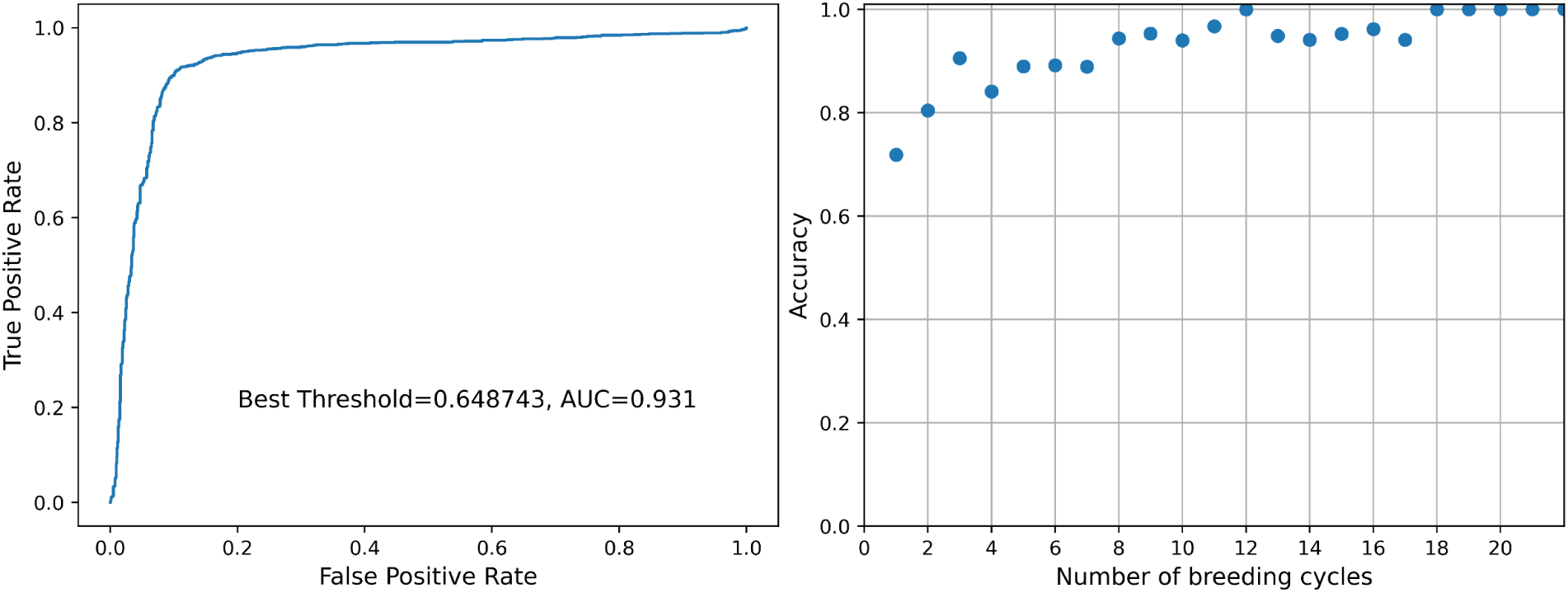
A) Receiver Operating Characteristic (ROC) curve of the sex probability for each individual given by the pooling of its lifetime sex probabilities against the molecular sexing. The Area Under the Curve (AUC)-ROC score and the best threshold are indicated in the figure. (B) Accuracy of sex classification according to the number of breeding cycles pooled together to determine the most probable sex of an individual. Higher number of breeding cycles leads to a better classification of the sex supporting the benefit of the pooling method.

**Figure S13:**
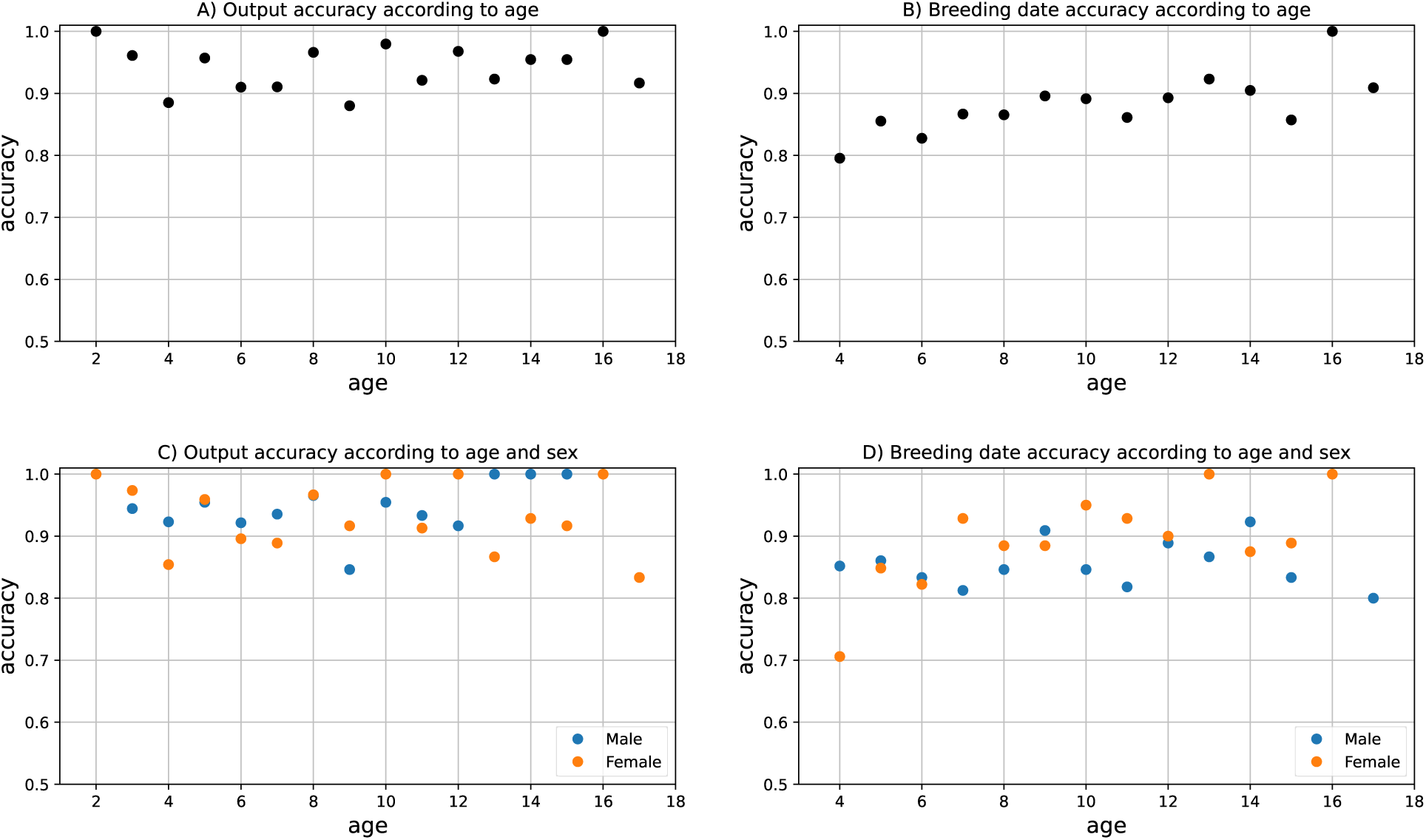
Age-specific (A, B, C, D) and sex-specific (C, D) accuracy metrics of output classification and breeding date determination. Output classification (Non-Breeder vs. Failure vs. Success) accuracy appeared stable according to age and sex. Age-specific accuracy metrics are given for age classes with sample sizes greater than 10 individuals. Accuracy of breeding date determination is slightly lower for young birds (breakpoint at 7 years old, from segmented regression analysis, ‘segmented’ R package).

